# TNF-α-Driven Systemic Inflammasome Hyperactivity Links Psoriatic Inflammation to monocyte inflammatory response and platelet activation

**DOI:** 10.1101/2025.11.24.690330

**Authors:** Deepti Verma, Freja Jeppsson, Florence Sjögren, Blanka Andersson, Cecilia Bivik Eding, Nada Katarina Kasic, Lavanya Moparthi, Charlotta Enerbäck

## Abstract

**Background:** Psoriasis is a chronic skin disease mediated by Th1 and Th17 immune responses and is classified as a systemic inflammatory disorder. Notably, psoriasis is an independent risk factor for myocardial infarction, stroke, and cardiovascular mortality, particularly in severe disease. However, the cellular mechanisms linking psoriasis to cardiovascular disease risk have not yet been identified. To address this gap, we investigated systemic markers of inflammasome signaling and innate immune activation in patients with psoriasis.

**Methods:** Whole blood was collected from 43 patients, including active psoriasis (mild-to-moderate disease) without clinical manifestations of atherosclerosis, inactive psoriasis (minimal disease activity), patients receiving anti-TNF-α therapy, and 19 BMI-matched healthy controls. Multiparametric spectral flow cytometry was performed to profile inflammasome signaling, and mass spectrometry-based proteomics was used to obtain unbiased phenotyping of circulating immune cells.

**Results:** Classical monocytes from patients with active psoriasis exhibited heightened NLRP3 protein expression and caspase-1 activity upon brief physiological stimulation, responses absent in inactive psoriasis and healthy controls. Mechanistically, active psoriasis demonstrated elevated plasma ATP and increased monocyte expression of P2X7R, a potent NLRP3 activator. TNF-α was identified as a key cytokine, selectively upregulating both P2X7R and NLRP3.

Baseline proteomics revealed enriched pathways for monocyte extravasation and cell adhesion, suggesting a pro-thrombotic state. Stimulation increased proteins linked to ROS and mitochondrial stress. Monocytes from active psoriasis exhibited increased baseline activation and, upon stimulation, enhanced monocyte-platelet aggregation, both of which were attenuated by inhibition of mitochondrial ROS. Importantly, anti-TNF therapy normalized ATP levels, P2X7R expression, inflammasome responsiveness, monocyte activation, and monocyte-platelet interactions, supporting the restoration of systemic immune homeostasis.

**Conclusions:** In patients with mild-to-moderate psoriasis, we demonstrate persistent systemic stress, resulting in inflammasome hyperreactivity and increased monocyte-platelet aggregation in response to minor perturbations in cellular homeostasis. Notably, TNF-α blockade restores these effects, providing mechanistic insight into how anti-TNF therapy reduces systemic inflammation and cardiovascular risk.

**What is already known about this topic?:**

- Psoriasis is a systemic, immune-mediated skin disease that is associated with an increased risk of cardiovascular disease (CVD), particularly in severe disease.
- The NLRP3 inflammasome, an innate immune sensor, has been implicated in the pathogenesis of CVD.
- Anti-TNF therapy, which is effective in treating psoriasis, is proposed to reduce CVD risk, but the underlying mechanisms remain unclear.

**What does this study add?:**

- Patients with mild-to-moderate psoriasis without clinical manifestations of atherosclerosis demonstrate elevated plasma ATP levels, increased monocyte P2X7 receptor expression, and enriched pathways for monocyte activation and extravasation, suggesting persistent systemic stress.
- Monocytes from these patients display ROS-dependent hyperactivation of the NLRP3 inflammasome and increased formation of monocyte-platelet aggregates in response to minor perturbations in cellular homeostasis.
- TNF-α was identified as a key cytokine, selectively upregulating both P2X7R and NLRP3.
- Anti-TNF therapy normalized these aberrant immune responses, suggesting a mechanism for its proposed cardioprotective effects.

**Novelty and significance:** This study provides evidence that patients with mild to moderate psoriasis, without clinical manifestations of atherosclerosis, exhibit concurrent elevations in plasma ATP levels, monocyte P2X7 receptor expression, inflammasome responsiveness, and monocyte-platelet aggregates: features increasingly associated with CVD risk. These findings suggest that dysregulated purinergic signaling may contribute to systemic immune activation in psoriasis. Importantly, anti-TNF therapy was associated with normalization of these parameters, pointing toward a potential immunomodulatory mechanism by which such treatment may help reduce CVD risk. These observations highlight a novel intersection between inflammation, purinergic signaling, and monocyte-platelet activation, which may contribute to the increased CVD risk in psoriasis.

## INTRODUCTION

Psoriasis is a chronic skin disease affecting around 2-3% of the worldwide population. It is characterized by hyperproliferation and abnormal differentiation of keratinocytes, as well as inflammation and angiogenesis in lesional skin. Specific cytokines, such as tumor necrosis factor (TNF)-α, interleukin (IL)-17, and IL-23, initiate and sustain the inflammation, and the blockade of these cytokines has been shown to be very effective as a treatment for severe psoriasis ^1^.

The systemic nature of psoriasis is increasingly being recognized ^2^. Psoriasis is linked to an increased risk of systemic comorbidities, including psoriatic arthritis, metabolic disorders, including obesity, and a pronounced risk for cardiovascular disease (CVD) ^2, 3^. While psoriasis is associated with an increased prevalence of traditional CVD risk factors, it is identified as an independent risk factor for CVD, with a higher estimated risk compared to individuals with type 2 diabetes ^4, 5^. Consequently, there is an increased risk of myocardial infarction, stroke, cardiovascular mortality, and reduced life expectancy (5 years), particularly in severe disease ^4–6^. The concept “psoriatic march” refers to the progression from chronic skin inflammation to systemic immune activation, ultimately leading to metabolic and vascular comorbidities ^7^.

The inflammasomes form the central hub of innate immune regulation. Members of the NOD-like receptor (NLR) family of proteins, including NLR family pyrin domain containing 3 (NLRP3), assemble into multiprotein complexes (termed ‘inflammasomes’) upon sensing danger signals^8^. The NLRP3 inflammasome is tightly controlled through a two-step process, where the first step, known as priming, involves the transcriptional upregulation of *NLRP3* and *pro-IL-1β*. The second step is activation of caspase 1, which cleaves pro-caspase-1 into its mature form, thereby releasing the proinflammatory cytokines IL-1β and IL-18 ^8^. The mechanosensitive purinergic P2X7 receptor is a crucial mediator in NLRP3 inflammasome activation ^9^. These receptors are ion channels that respond to extracellular ATP, which can be released during cellular stress, inflammation, or mechanical stimulation. Once activated, these receptors facilitate Ca2+ influx and K+ efflux, which are essential for NLRP3 inflammasome assembly and activation^9, 10^.

The activation of the NLRP3 inflammasome is increasingly recognized as a critical factor in the development and progression of atherosclerosis^11–13^. The Canakinumab Anti-Inflammatory Thrombosis Outcomes Study (CANTOS) revealed a lower rate of recurrent cardiovascular events in patients treated with the anti-IL-1β antibody canakinumab ^14^, thereby directly linking inflammasome-driven inflammation to CVD.

We have previously shown an association between single-nucleotide polymorphism in *NLRP3* and psoriasis susceptibility ^15^. More recently, we reported that monocytes from patients with psoriasis exhibit enhanced caspase-1 response to lipopolysaccharide (LPS) compared to healthy controls, suggesting a hyperresponsive inflammasome state. TNF-α was found to selectively activate the NLRP3 inflammasome without requiring a priming signal ^16^.

Although systemic inflammation in psoriasis is anticipated to increase CVD risk, the underlying cellular mechanisms remain incompletely understood. Here, we demonstrate that intrinsic stress lowers the activation threshold of psoriatic monocytes, leading to inflammasome hyperactivity and the formation of monocyte-platelet aggregates in response to minor perturbations in cellular homeostasis. Furthermore, we identify TNF-α as a regulator of systemic inflammasome function in patients with psoriasis.

## METHODS

### Study subjects

Patients diagnosed with psoriasis vulgaris were recruited from the Department of Dermatology, Linköping University Hospital, Linköping, Sweden. Disease severity was assessed using the Psoriasis Area and Severity Index (PASI). Patients with mild to moderate PASI scores were classified as having active disease, whereas those with minimal disease activity were classified as having inactive disease ^17^. Laboratory parameters, including total cholesterol, high- and low-density lipoprotein (HDL, LDL), triglycerides, HbA1c, and high-sensitivity (hs) CRP were determined. White blood cell (WBC)-derived inflammatory indices, including monocyte-to HDL ratio (MHR), monocyte-to-lymphocyte ratio (MLR), neutrophil-to-lymphocyte ratio (NLR), and platelet-to-lymphocyte ratio (PLR), were calculated.

Exclusion criteria included biological treatment within the past 4 months, UVB therapy exceeding 3 sessions, pregnancy, malignancy within the last 5 years, active infection within the past 72 hours, and treatments known to affect the studied markers.

Patients undergoing anti-TNF-α therapy had received the treatment for at least one year. Written informed consent was obtained from all participants. The study was approved by the Local Ethics Committee and conducted in accordance with the principles of the Declaration of Helsinki.

### Blood sample collection and processing

Our studies were performed by promptly processing freshly isolated blood cells to circumvent the confounding effects associated with prolonged cell cultures, which may modify sensitive cellular responses. Peripheral blood was collected from study participants in EDTA-containing tubes. Plasma was isolated by centrifugation at 2000 x g for 10 minutes at room temperature and stored at -70°C for subsequent analyses. The remaining blood samples were immediately processed for downstream analysis as described in the later sections.

All samples were analyzed at room temperature (20-22°C) and after exposure to 37°C for 10 minutes. The response at 37°C was quantified as the fold change relative to the value at room temperature. Mitoquinone (MitoQ, 2.45 µM; Cayman chemicals), bzATP (50-100 µM; Sigma-Aldrich), and TNF-α (10ng/ml; R&D systems) were used.

### Immunostaining and flow cytometry

#### Panel for the detection of NLRP3 (Panel 1, 9-fluorochrome panel)

Following treatment of whole blood with Brefeldin A (GolgiPlug, BD Biosciences), the red blood cells were lysed using BD Pharm Lyse (BD Biosciences). Cells were incubated with extracellular antibodies (Supplementary Table S1) for 30 minutes at 4°C. After washing, cells were fixed with 2% paraformaldehyde at 4°C overnight. Following permeabilization with Cytofix/Cytoperm (BD Biosciences), samples were incubated for 30 minutes with anti-NLRP3-PE (R&D Systems) antibodies (Supplementary Table S1). Monocytes were identified based on their characteristic forward scatter (FSC) and side scatter (SSC) properties. A minimum of 10,000 events within the monocyte gate were acquired for each sample.

#### Detection of active caspase-1, P2X7, CD40 and CD41(Panel 2, 14- fluorochrome panel)

Peripheral blood mononuclear cells (PBMCs) were isolated using Ficoll-Paque Plus gradient centrifugation (SerumWerk Bernburg) in SepMate PBMC isolation tubes (STEMCELL Technologies) according to the manufacturer’s instructions. Cells were incubated with extracellular antibodies, including P2X7, CD40, and CD41 (Supplementary Table S1) for 30 minutes on ice. After washing, cells were incubated with a Fluorescence inhibitor of caspase-1 (FLICA) reagent (ImmunoChemistry Technologies) for 30 minutes at 37°C. Data were acquired on a spectral flow cytometer. Monocytes were identified based on their characteristic forward scatter (FSC) and side scatter (SSC) properties. A minimum of 50,000 events within the monocyte gate were acquired for each sample. Monocyte platelet aggregates were identified based on their scatter properties and co-expression of CD14 and CD41. Antibodies were titrated, and gates were set based on negative controls, single-staining, and fluorescence minus one sample. Data were acquired on a 3L Cytek Aurora spectral flow cytometer (Cytek Biosciences). Flow cytometric analysis was performed using Kaluza version 2.1 software. Data are presented as median fluorescence intensity (MFI).

### ATP quantification

Extracellular ATP levels were quantified in plasma samples and supernatants from PBMCs. For collecting the cell culture supernatants, 2 × 10^6 PBMCs were resuspended in 100 μL PBS and incubated at room temperature (20-22°C) or 37°C for 10 or 60 minutes. Following incubation, samples were centrifuged at 400 × g for 5 minutes. Supernatants were collected and stored at -70°C until analysis. ATP concentrations were measured using the ATP Determination Kit (Molecular Probes, Invitrogen), following the manufacturer’s protocol. Luminescence was recorded using a microplate reader (SpectraMax Gemini Fluorometer, Molecular Devices). ATP concentrations were calculated using a standard curve ranging from 0.01µM to 35 µM. Each sample was measured in duplicate.

### RNA extraction and quantitative Real-Time (RT) PCR

PBMCs were lysed in RLT buffer (Qiagen) and stored at -80°C. RNA was purified using the RNeasy micro plus kit (Qiagen) and quantified using the Nanodrop spectrophotometer (Thermo Fisher Scientific™), following the manufactureŕs instructions. cDNA synthesis was conducted with the Maxima First Strand cDNA Synthesis Kit for RT-qPCR (Thermo Fisher Scientific™). Quantitative real-time PCR was performed in the QuantStudio™ 7 Flex Real-Time PCR System (Applied Biosystems™) using SYBR Green assay (Applied Biosystems™) to determine the expression of *IL-1β* and TaqMan assay (Applied Biosystems™) for *CD39* and *CD73*. Data collection was conducted in triplicate and normalized to *RPLP0* using the comparative Ct method 2^-ΔΔCt^ for calculating fold changes and 2^-ΔCt^ for baseline comparisons.

### Sample preparation for mass spectrometry

Peripheral venous blood was collected in EDTA tubes from healthy individuals and patients with psoriasis. Peripheral blood cells were pelleted from whole blood after lysing red blood cells using 1 x BD Pharm Lyse™ lysing solution (BD Biosciences). The cell pellets were incubated at the mentioned temperatures to prime the cells. The cell pellets were lysed using RIPA buffer (Thermo Fisher Scientific™) containing complete mini protease inhibitors (Thermo Fisher Scientific) and sonicated. Cell debris was removed by centrifuging at 13000 rpm for 10 minutes at 4°C. The protein concentrations were determined by bicinchoninic acid (BCA) assay (Thermo Fisher Scientific). Sample preparation was performed using the Filter Aided Sample Preparation (FASP) method ^18^. For each sample, 20-100 µg of total protein was used and reduced with 10 mM dithiothreitol (DTT) at 95 °C for 5 minutes on a thermomixer. After cooling, the samples were denatured with urea buffer (8 M urea in 0.1 M Tris-HCl, pH 8.5) in a total volume of 200 µl and transferred to the 10 kDa filter unit (Millipore) and centrifuged at 12000 g for 20 minutes. The denatured proteins were alkylated with 100 µl of 50 mM Iodoacetamide in the urea buffer in the filter unit and incubated at room temperature in the dark for 30 minutes and then centrifuged at 12000 g for 20 minutes. The filter units were further washed three times with the urea buffer. Then, the samples were subsequently washed again with Tris buffer (0.1 M Tris-HCl, pH 8.5) to remove urea before trypsin digestion. The samples were digested on filter units with mass spectrometry grade Pierce™ Trypsin/Lys-C Protease Mix (Thermo Fisher Scientific) in a 1:50 (enzyme: protein) ratio at 37 °C overnight. Digested samples were desalted using C18 spin columns (Thermo Fisher Scientific), lyophilized under vacuum, and stored at -20 °C.

### Liquid chromatography and mass spectrometry

The desalted and dried peptide samples were reconstituted in 0.1% Formic acid (FA) in ultra-pure Milli-Q water, and the concentration was measured using a Nanodrop (Thermo Fisher Scientific). The samples were analyzed using a nanoElute 2 LC system connected with a timsTOF HT mass spectrometer via the CaptiveSpray 2 source (Bruker). The peptides were separated by a 25 cm C18 column (150 μm inner diameter, 1.5 μm particle size, PepSep, Bruker Daltonics) with a gradient of 2-17 % solvent B (0.1% FA in acetonitrile (ACN)) in 25 min, 17-25% B in 5 minutes, 25-37% B in 5 minutes, 37-95% B in 10 minutes at a flow rate of 400 nl/minutes. The MS data were acquired using the dia-PASEF method. The capillary voltage was set to 1600 V. The MS and MS/MS spectra were acquired from 100-1700 m/z. The ion mobility was scanned from 0.85 to 1.3 Vs/cm2. The ramp time was set to 100ms. The collision energy was ramped linearly as a function of the mobility from 59 eV at 1/K0= 1.6 Vs/cm2 to 20 eV at 1/K0= 0.6 Vs/cm2.

### Protein identification and quantification

The raw DIA data files were converted into HTRMS files using the HTRMS Converter (version 18.7, Biognosys). These HTRMS files were processed using the directDIA+(Deep) workflow in Spectronaut proteomics software (version 19.1.240724.62635, Biognosys). The data files were searched against the UniProtKB-proteome UP000005640 (Taxonomy: Homo sapiens, release: 2024-04-02, Entries: 82493). The default BGS Factory Settings were used for search and analysis with the following modifications: Trypsin/P and LysC/P were used as specific enzymes allowing for a maximum of two missed cleavages; Peptide length from 5 to 52; Carbamidomethylation of cysteines was set as fixed modification whereas Oxidation of methionine and N-acetylation were set as variable modifications; The identifications were filtered at FDRs of 0.01% on PSM, peptide and protein level; Excluded single-hit proteins; Missing values were imputed using global imputation strategy and cross-run normalization was enabled.

Gene ontology enrichment analysis was performed using the Gene Ontology (GO) resource (geneontology.org)^19, 20^ powered by PANTHER (release 19). The Fisher’s exact test was conducted, and false discovery rate (FDR)-corrected values were used.

### Statistics

Data analysis was performed in GraphPad Prism version 10.3.1 (GraphPad Software Inc., San Diego, CA, USA). Normality was assessed using a combination of visual inspection (Q–Q plots) and statistical tests (Shapiro–Wilk test and/or Kolmogorov–Smirnov test). Variables were considered normally distributed when the p-value exceeded 0.05 and visual plots indicated symmetry. p-values were calculated using ANOVA or Kruskal-Wallis test for continuous variables, and Chi-square test or Fisheŕs exact test for categorical variables, where appropriate. Post hoc analyses were performed (Šídák’s, Dunn, or Tukey). Correlations were determined using Pearson correlation coefficients.

## RESULTS

A total of 43 patients were enrolled: 16 with active psoriasis, 13 with inactive psoriasis, 14 receiving anti-TNF therapy, and 19 healthy controls. The mean duration of psoriasis was over 20 years in all patient groups. Baseline characteristics of all groups are summarized in Table 1. The mean PASI was 8.4 ± 4.36 in patients with active psoriasis and 1.75 ± 0.62 in those with inactive psoriasis. Active psoriasis patients had a comparable age to healthy controls (51.5 ± 12.8 vs. 58.8 ± 7 years) and showed no previous signs or symptoms of atherosclerosis. BMI was matched across all groups (active psoriasis 27.3 ± 4.9, inactive psoriasis 26.8± 3.3, anti-TNF-treated psoriasis 27.4± 3.7, and controls 24.9 ± 3.8 kg/m²).

**Table 1:**
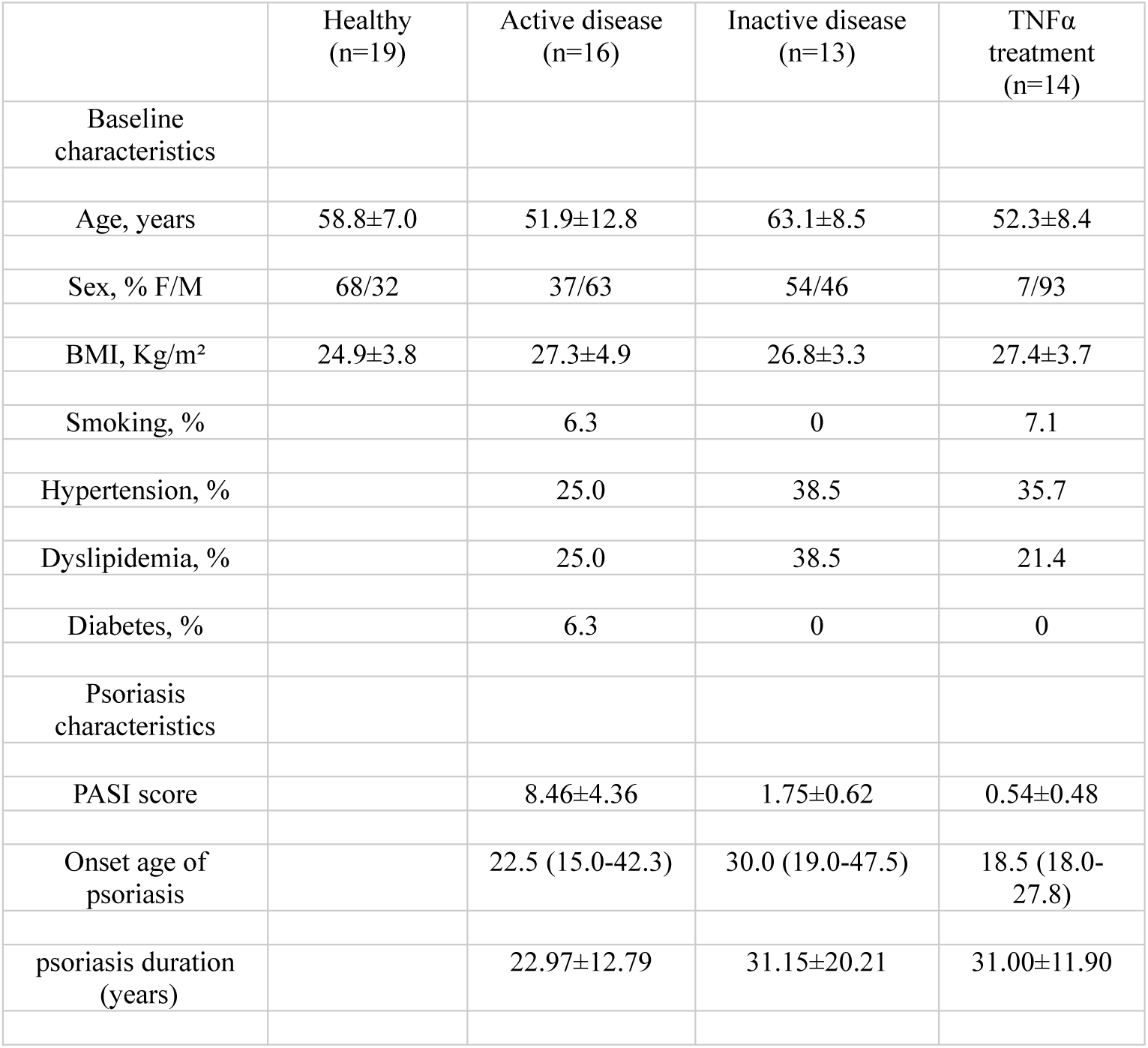
Baseline characteristics of subjects in the study Data are presented as mean (SD) for parametric variables and median (IQR) for non-parametric variables. Among inactive psoriasis patients, one had a previous stroke, and another had a previous myocardial infarction. No other signs of atherosclerosis were reported in the other patients. Abbreviations: BMI, body mass index; F/M, female/male; PASI, psoriasis area severity index.

No significant differences in lipid, metabolic, or inflammatory markers were observed between the patient groups (Table 2). All patient groups had median hsCRP levels below 2mg/L, which indicates moderate vascular risk^21^. Additionally, WBC-derived inflammatory indices (MHR, MLR, NLR, and PLR) were within normal range.

**Table 2:** Baseline levels of metabolic and inflammatory markers Data are presented as mean (SD) for parametric variables and median (IQR) for non-parametric variables. Patients treated with statins are excluded from lipid profile and MHR. Abbreviations: HbA1c, hemoglobin A1c; HDL, high-density lipoprotein; hsCRP, high-sensitivity CRP; LDL, low-density lipoprotein; MHR, monocyte-to-HDL ratio; MLR, monocyte-to-lymphocyte ratio; NLR, neutrophil-to-lymphocyte ratio; PLR; platelet-to-lymphocyte ratio.

| | Active disease | Inactive disease | TNFα treatment |
| --- | --- | --- | --- |
| Metabolic markers |  |  |  |
| Total cholesterol, mmol/L | 5.26±1.01<br>[n=8] | 6.35±1.05<br>[n=8] | 5.44±1.32<br>[n=10] |
| HDL cholesterol, mmol/L | 1.55 (1.40-1.95)<br>[n=8] | 1.60 (1.35-2.08)<br>[n=8] | 1.15 (1.00-1.50)<br>[n=10] |
| LDL cholesterol, mmol/L | 3.66±1.11<br>[n=8] | 4.69±1.26<br>[n=8] | 3.65±1.20<br>[n=10] |
| HbA1c mmol/mol | 36.00±3.86<br>[n=10] | 38.55±3.48<br>[n=11] | 36.2±5.03<br>[n=10] |
| Inflammatory markers |  |  |  |
| LPK × 10<sup>9</sup>celler/L | 6.40 (4.93-7.60)<br>[n=12] | 6.30 (4.80-7.30)<br>[n=11] | 6.10 (4.83-7.53)<br>[n=14] |
| TPK × 10<sup>9</sup>celler/L | 240.30±39.09<br>[n=8] | 229.00±53.40<br>[n=11] | 246.10±34.27<br>[n=9] |
| hsCRP mg/L | 1.20 (0.73-5.00)<br>[n=8] | 1.60 (0.80-1.90)<br>[n=11] | 1.55 (0.65-2.55)<br>[n=8] |
| MHR | 0.34 (0.24-0.69)<br>[n=8] | 0.27 (0.21-0.39)<br>[n=8] | 0.53 (0.28-0.67)<br>[n=9] |
| MLR | 0.29 (0.25-0.36)<br>[n=11] | 0.28 (0.24-0.43)<br>[n=11] | 0.25 (0.21-0.30)<br>[n=9] |
| NLR | 1.83 (1.76-2.88)<br>[n=11] | 1.95 (1.63-2.75)<br>[n=11] | 1.36 (1.17-1.91)<br>[n=9] |
| PLR | 159.10±71.89<br>[n=7] | 132.30±39.28<br>[n=11] | 109.20±29.69<br>[n=9] |

### Monocytes from active psoriasis display an increased inflammasome reactivity to physiological stimuli

Using an extended flow cytometry panel, modified from Ong et al. ^22^, we found no significant differences in the percentages of circulating monocyte subpopulations between psoriasis patients and controls. We observed predominant NLRP3 expression in the classical monocyte population, as supported by Giral et al. ^23^, and therefore focused our subsequent analyses on this monocyte subset.

We explored whether there are differences in the functional and activation states of psoriatic monocytes, with a particular focus on the NLRP3 inflammasome. We observed an unexpected increase in NLRP3 protein expression in classical monocytes from patients with active psoriasis when the PBMC samples were exposed to a short physiological stimulus by transitioning them from room temperature to 37°C for 10 minutes (Figure 1A). In addition, caspase 1, which reflects inflammasome activity, was also significantly enhanced (Figure 1B). The activation of the inflammasome was also accompanied by significantly increased pro-*IL-1β* mRNA levels (Figure 1C). These responses were not observed in inactive psoriasis or in healthy controls.

**Figure 1.**
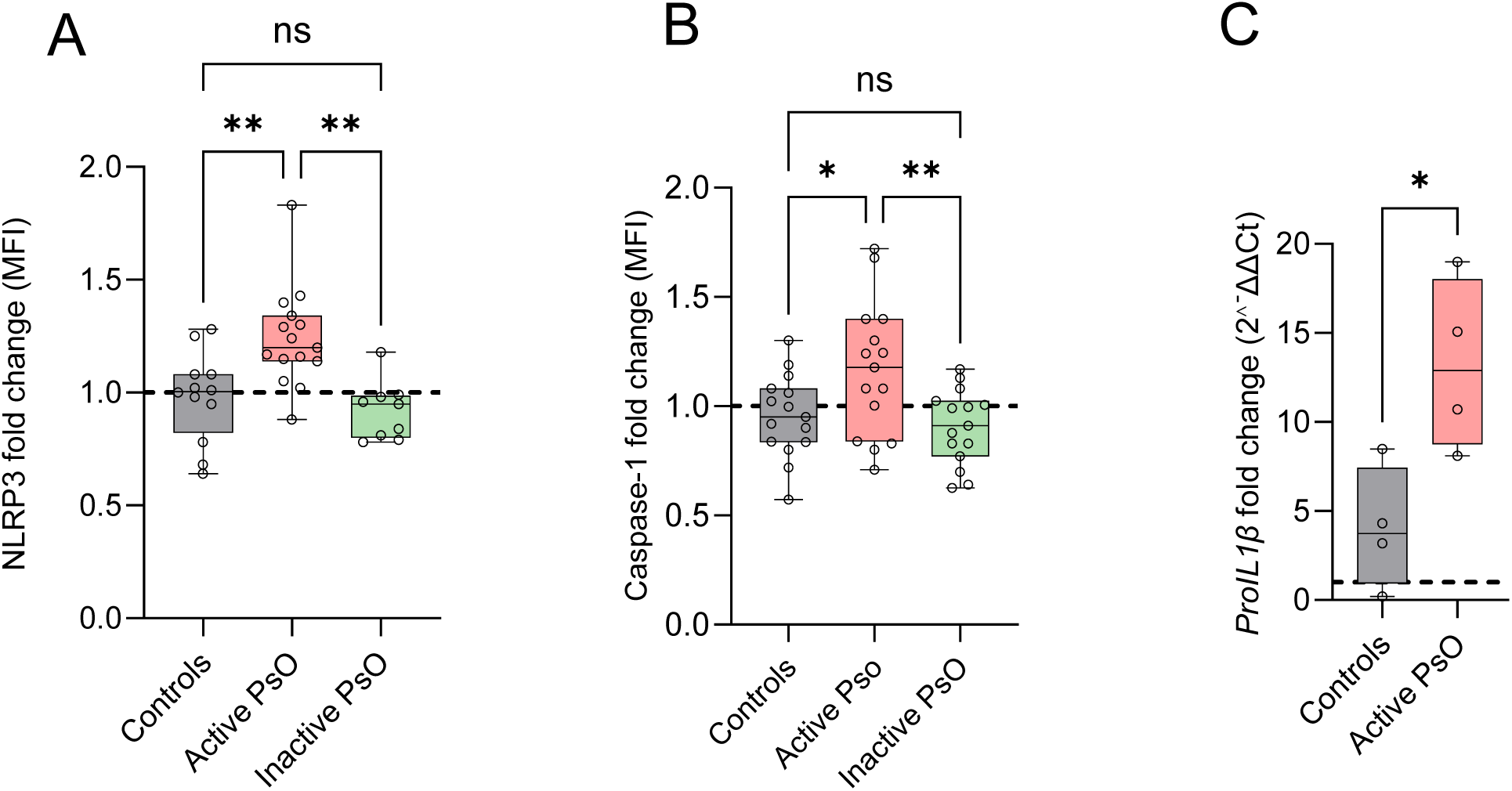
Increased inflammasome activity in classical monocytes from patients with active psoriasis (PsO) in response to physiological stimulation. (A) Ratio of median fluorescence intensity (MFI) for NLRP3 expression at 37 °C versus room temperature for 10 minutes, measured by spectral flow cytometry (n = 9–16 per group). (B) Ratio of caspase-1 MFI at 37 °C versus room temperature for 10 minutes, measured by spectral flow cytometry (n = 15–17 per group). (C) Ratio of pro-IL-1β mRNA expression at 37 °C versus room temperature for 10 minutes, determined by quantitative PCR (n = 4 per group). *p <0.05, ** p < 0.01, *** p<0.001, ns = non-significant, one-way ANOVA was used for comparisons involving more than two groups, and Šídák’s post hoc test was performed for multiple comparisons. Student’s t-test was used for comparisons between two groups.

Notably, the basal levels of these proteins were similar among the groups (Supplementary Figure S1). Our findings suggest that monocytes from active psoriasis are in a heightened readiness, requiring minimal additional activation to trigger a full inflammatory response.

### Enhanced constitutive P2X7 receptor expression in active psoriasis

To investigate the underlying mechanism of hyperreactivity in psoriatic monocytes, we focused on the extracellular ATP receptor P2X7, a well-characterized and potent activator of the NLRP3 inflammasome ^10^. P2X7R is temperature-sensitive, and enhanced activation has been observed upon transitioning samples from 4°C to 37°C ^24^.

We found significantly higher constitutive P2X7R levels in patients with active psoriasis (Figure 2A) than in healthy controls, further supporting the primed state of psoriatic monocytes. Importantly, no differences in TOPRO-3 dye uptake (Figure 2B) were observed, suggesting that the elevated P2X7R surface expression was not accompanied by large pore formation characteristic of osmotic cell lysis (pyroptosis).

**Figure 2.**
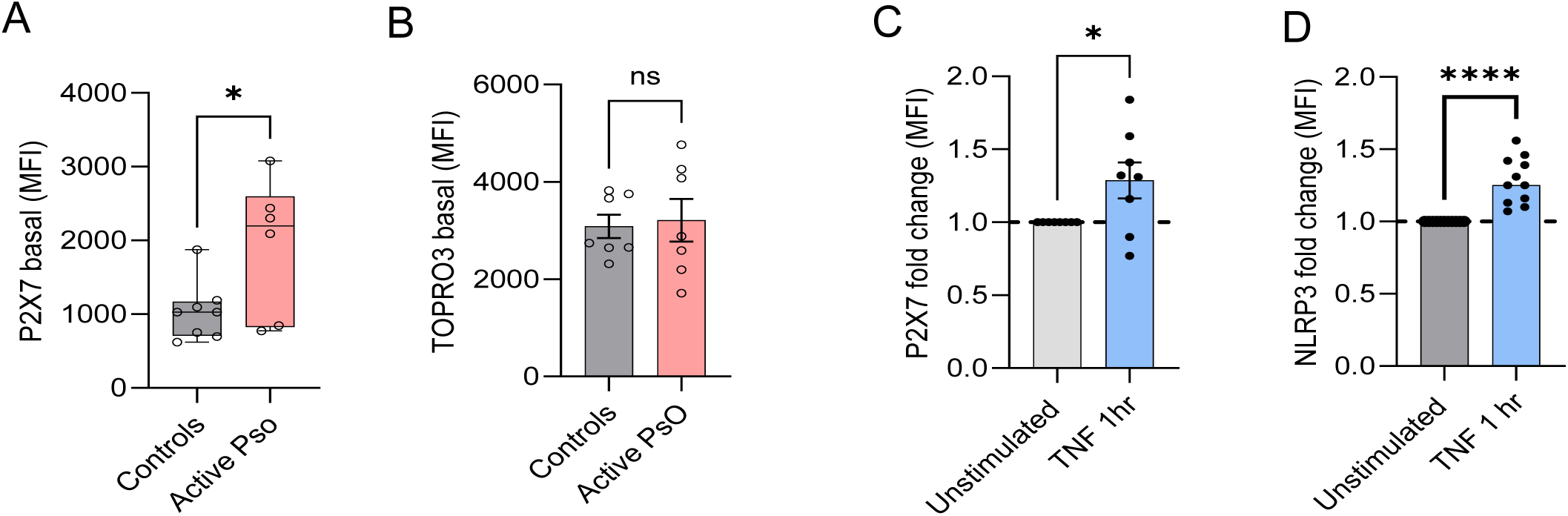
Increased basal P2X7 surface expression in monocytes from patients with active psoriasis (PsO). (A) Baseline P2X7 receptor surface expression (MFI) on classical monocytes from individuals with active PsO, measured by spectral flow cytometry (n = 6–8 per group). (B) Baseline TOPRO3 dye uptake (MFI) by classical monocytes, measured by spectral flow cytometry (n=7 per group). (C) Effect of 1 hr in vitro treatment with TNF-α (10ng/ml) on P2X7 (MFI) in monocytes (n=8). (D) Effect of 1 hr in vitro treatment with TNF-α on NLRP3 (MFI) in monocytes (n=11 per group). Data are presented as mean ±SEM. *p <0.05, **** p<0.0001. ns = non-significant, Student’s t-test was used for comparisons between two groups.

We previously demonstrated that among systemically elevated cytokines in psoriasis, including IL-17, IL-23, and IFN-γ, only TNF-α specifically induces NLRP3 transcription ^16^. We now demonstrate that TNF-α treatment significantly enhances surface expression of the P2X7 receptor (Figure 2C) and upregulates NLRP3 (Figure 2D) at the protein level. These data underscore the critical role of TNF-α in priming monocytes through modulation of key inflammasome components.

### Increased plasma ATP levels in active psoriasis

ATP, the primary agonist of P2X7R, triggers a cascade of events, resulting in NLRP3 inflammasome assembly and caspase-1 activation ^10^. The increased expression of P2X7R in psoriasis raises the hypothesis of elevated plasma levels of its ligand, ATP. Using a bioluminescence assay, we observed substantially higher plasma ATP levels in active psoriasis (mean 13.66 µM ± 6.3) compared to inactive psoriasis (5.3µM ± 3.9) and healthy controls (4.9µM ± 1.29 Figure 3A). The co-elevation of ATP, a well-established marker of cellular stress, and P2X7R indicates an enhanced systemic stress response in active psoriasis. Notably, there were no statistically significant differences in ATP levels between inactive psoriasis and healthy controls. Furthermore, plasma ATP levels correlated with disease severity (Pearson correlation = 0.51, *p* = 0.02) and with caspase-1 activity in response to physiological stimulation (Pearson correlation = 0.56, *p* = 0.02) (Figure 3B&C).

**Figure 3.**
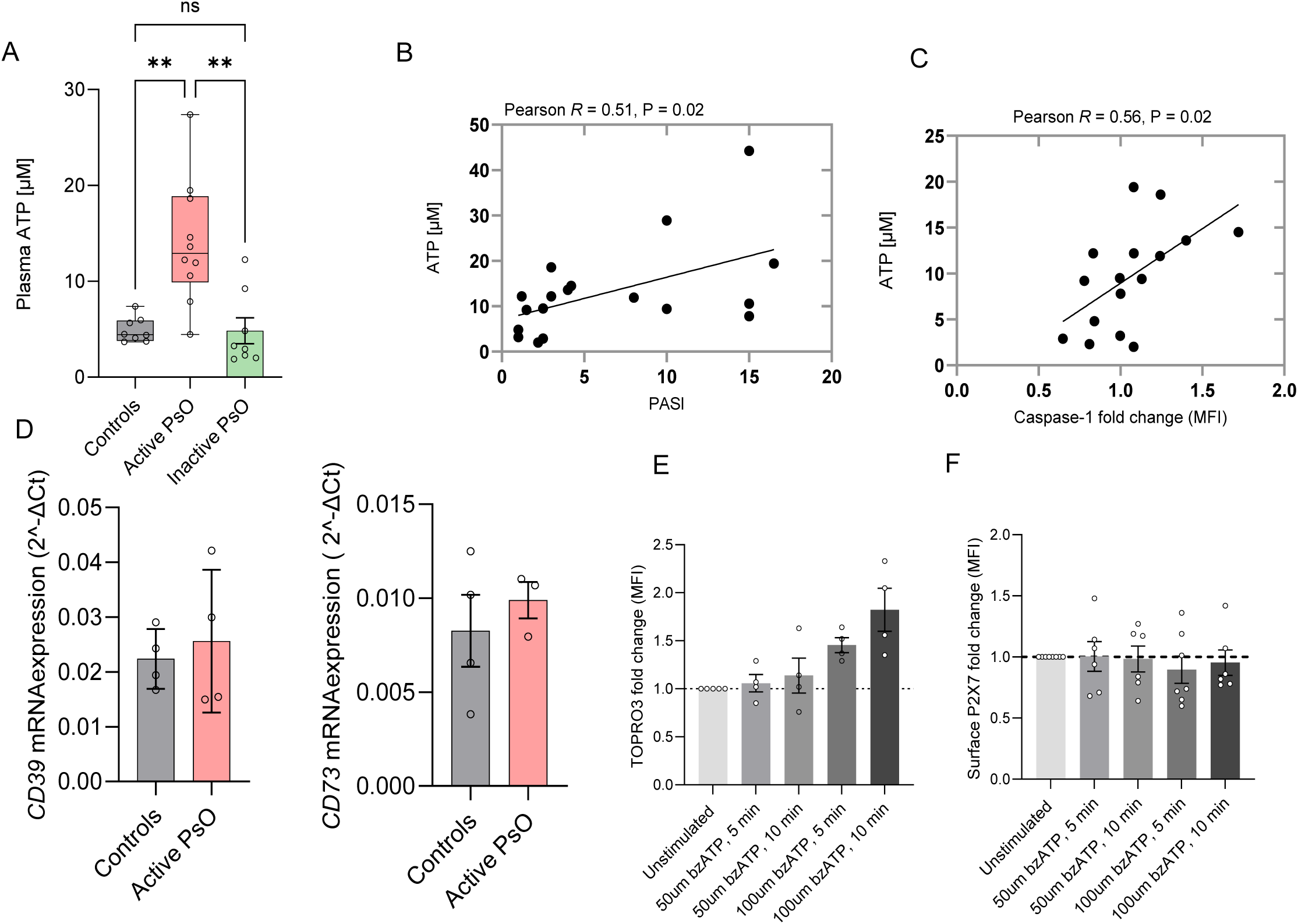
Increased plasma ATP levels in patients with active psoriasis (PsO). (A) Plasma ATP concentration measured by bioluminescence assay (n = 8–11 per group). (B) Correlation between plasma ATP (µM) and Psoriasis Area and Severity Index (PASI), assessed using Pearson’s correlation coefficient (n = 18). (C) Correlation between plasma ATP and the ratio of caspase-1 response at 37 °C versus room temperature, assessed using Pearson’s correlation coefficient (n = 16). (D) Baseline *CD39* and *CD79* mRNA levels were determined by quantitative PCR (n = 4 per group). (E) Effect of a 5 and 10-minute treatment with bzATP (50 µM and 100 µM) on TO-PRO-3 dye uptake in monocytes and (F) on P2X7 surface expression on monocytes, measured by spectral flow cytometry (n = 4–6 per group). Data are presented as mean ±SEM. ** p < 0.01, ns = non-significant, one-way ANOVA was used for comparisons involving more than two groups, and Šídák’s post hoc test was performed for multiple comparisons. Student’s t-test was used for comparisons between two groups.

The half-life of extracellular ATP is short, since it is rapidly hydrolyzed by the ectonucleotidases CD39 and CD73 ^25^. We found no changes in the constitutive mRNA expression levels of *CD39* and *CD73* in PBMCs from active PsO, suggesting that the increased plasma ATP is not due to dysregulated ATP hydrolysis. (Figure 3D).

To evaluate the effect of ATP on P2X7 surface expression, cells were stimulated with the selective agonist bzATP. Increasing concentrations of bzATP resulted in a dose-dependent increase in P2X7-mediated pore formation, as measured by TOPRO3 uptake (Figure 3E). Notably, this functional response was observed in the absence of detectable changes in the abundance of cell surface receptors (Figure 3F). Moreover, no synergistic effect was observed when TNF-α and ATP were combined (data not shown), indicating that TNF-α contributes independently to receptor upregulation. These findings point out the distinct regulatory roles of ATP and TNF-α in the modulation of the P2X7R.

### Proteomic analysis reveals primed immune cells and augmented ROS pathway activation in active psoriasis

To obtain an unbiased global proteomic profile of circulating immune cells following stimulation, we performed high-resolution mass spectrometry on erythrocyte-depleted blood cells from patients with active psoriasis and healthy controls. In response to stimulation, active psoriasis samples demonstrated gene ontology (GO) enrichments in defense response, response to stress, immune system process, and response to external stimulus (Figure 4A), highlighting a primed, hyperreactive immune state that extends beyond inflammasome activation. In contrast, no pathway enrichments were detected in healthy controls, demonstrating that the stimulation was insufficient to activate immune cells from healthy individuals.

**Figure 4.**
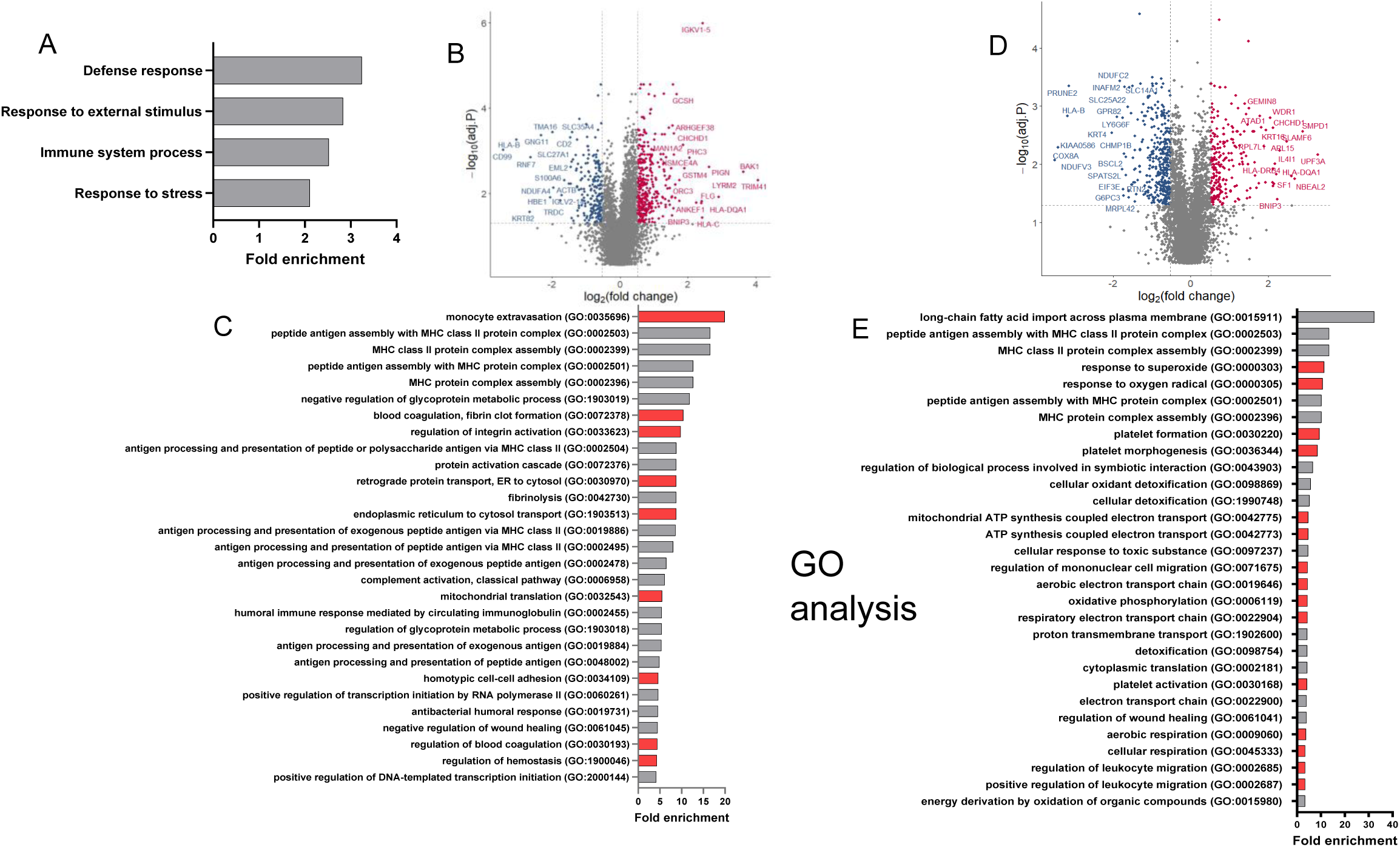
Increased response of primed immune cells to physiological stimuli in active psoriasis(PsO). (A) Gene ontology (GO) analysis of biological processes following stimulation in immune cells from patients with active PsO, n=6. (B) Volcano plot displaying differential gene expression in active PsO vs controls at baseline. (C) GO analysis of biological processes comparing active PsO and controls at baseline. Volcano plot and GO analysis following stimulation (D &E), comparing active PsO and controls, n=6 per group. Red and blue colors in volcano plots (B&D) indicate differentially over- or under-expressed proteins, respectively, in active PsO vs. controls. Red-colored bars in the GO analysis indicate pathways associated with monocyte or platelet function that are relevant for this study.

To investigate inherent differences between active psoriasis patients and healthy controls, we compared their proteomic profiles (active psoriasis vs controls) at baseline and after stimulation.

At baseline (Figure 4 B&C), the most significantly enriched gene ontology (GO) term was ‘monocyte extravasation’, indicating active monocytes equipped for tissue recruitment. This was further supported by enrichments in cell-cell adhesion, coagulation, and hemostasis, further supporting a pro-inflammatory and pro-thrombotic basal state in active psoriasis. In addition, significant enrichments were detected in pathways of mitochondrial translation and endoplasmic reticulum, supporting a constitutively active or primed state in active psoriasis.

Upon stimulation (Figure 4 D&E), we observed significant enrichments of pathways linked to reactive oxygen species (ROS) production, mitochondrial metabolism, energy generation, oxidative stress, and platelet activation signaling. These findings collectively suggest increased monocyte activation and oxidative stress in active psoriasis, which may mechanistically contribute to vascular inflammation and cardiovascular risk.

### Increased monocyte activation and monocyte-platelet aggregate (MPA) formation in psoriasis

Platelet activation and interactions with immune cells, especially monocytes, play a pivotal role in the pathogenesis of cardiovascular disease ^26^. In line with our proteomic data, we found that monocytes from patients with active psoriasis exhibited elevated baseline CD40 expression compared to healthy controls (Figure 5A), which reflects an active monocyte phenotype. Next, we assessed the expression of CD41, a platelet-specific marker to detect monocyte-platelet aggregates, an established marker of in vivo platelet activation linked to coronary artery disease ^27^. We found no significant difference in basal CD41 expression on monocytes between the groups (data not shown. Upon stimulation, however, classical monocytes from patients with active psoriasis, unlike those from controls, exhibited rapidly increased MPA formation (Figure 5B). Inhibition of mitochondrial ROS (mtROS) with MitoQ significantly reduced monocyte activation (Figure 5C), MPA formation (Figure 5D), and caspase-1 activation (Figure 5E), underscoring the central mechanistic role of mitochondrial stress in both platelet and inflammasome activation.

**Figure 5.**
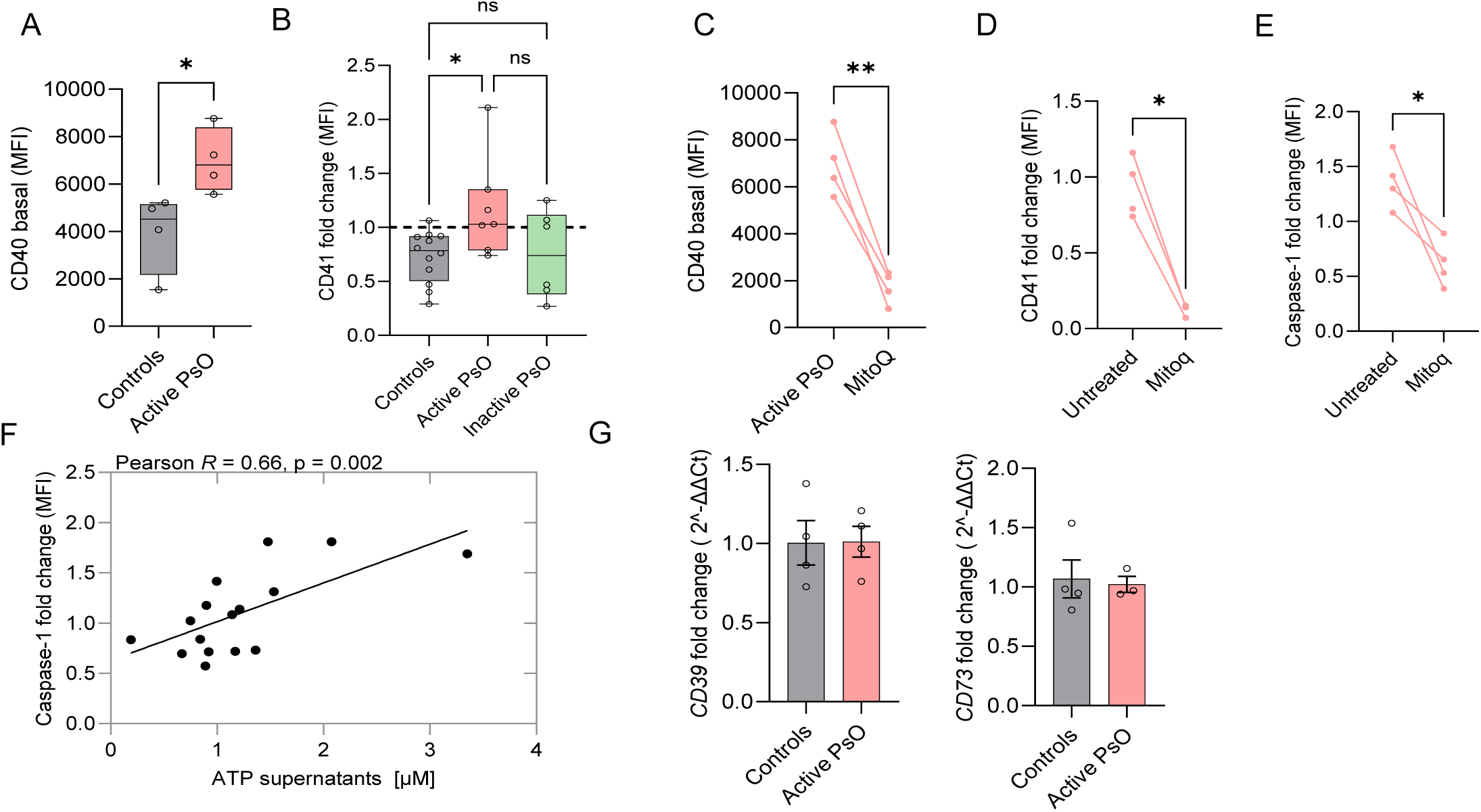
Increased baseline monocyte activation and platelet adhesion to monocytes in patients with active psoriasis (PsO). (A) Baseline CD40 expression (MFI) on classical monocytes, measured by spectral flow cytometry (n=4-5 per group). (B) Ratio of CD41 expression (MFI) on classical monocytes at 37 °C versus room temperature for 10 minutes, measured by spectral flow cytometry (n=6-12 per group. (C) CD40 expression (MFI) upon treatment with MitoQ (2.5μM), (n=4). (D) CD41 fold change upon treatment with MitoQ (2.5 μM), (n=4 per group). (E) Caspase-1 fold change (MFI) upon treatment with MitoQ (2.5μM), n=4. (F) Correlation between caspase-1 fold change (MFI) and ATP release (μM) into supernatants at 37 °C, assessed using Pearson’s correlation coefficient (n= 16). (G) Fold change mRNA levels in *CD39/CD73* as determined by qPCR. Data are presented as mean ±SEM. *p <0.05, ** p < 0.01, ns = non-significant. One-way ANOVA was used for comparisons involving more than two groups, and Šídák’s post hoc test was performed for multiple comparisons. Student’s t-test was used for comparisons between two groups.

Since the proteomic data indicated increased ATP synthesis, we quantified ATP release following stimulation and found that ATP strongly correlated with caspase-1 activity (Pearson R = 0.66, p = 0.002, Figure 5F). As anticipated, mRNA levels of *CD39* and *CD73* remained comparable between groups (Figure 5G) upon stimulation, indicating a similar ATP degradation.

In conclusion, monocytes from active psoriasis exhibited increased baseline activation and enhanced monocyte-platelet aggregate formation after stimulation, both of which decreased with mitochondrial ROS inhibition.

### Anti-TNF treatment normalizes P2X7R expression and plasma ATP levels

TNF-α inhibition using neutralizing antibodies is highly effective in treating psoriatic skin lesions ^28^. Based on our findings that TNF-α upregulates P2X7R expression, we recruited patients who had been effectively treated with anti-TNF antibody therapy for at least one year.

We examined the expression of P2X7R on classical monocytes and determined the plasma ATP levels in this cohort. Interestingly, the P2X7R expression in these patients was similar to that observed in controls (Figure 6A). Furthermore, plasma ATP levels in these patients were normalized to those of healthy controls (Figure 6B).

**Figure 6.**
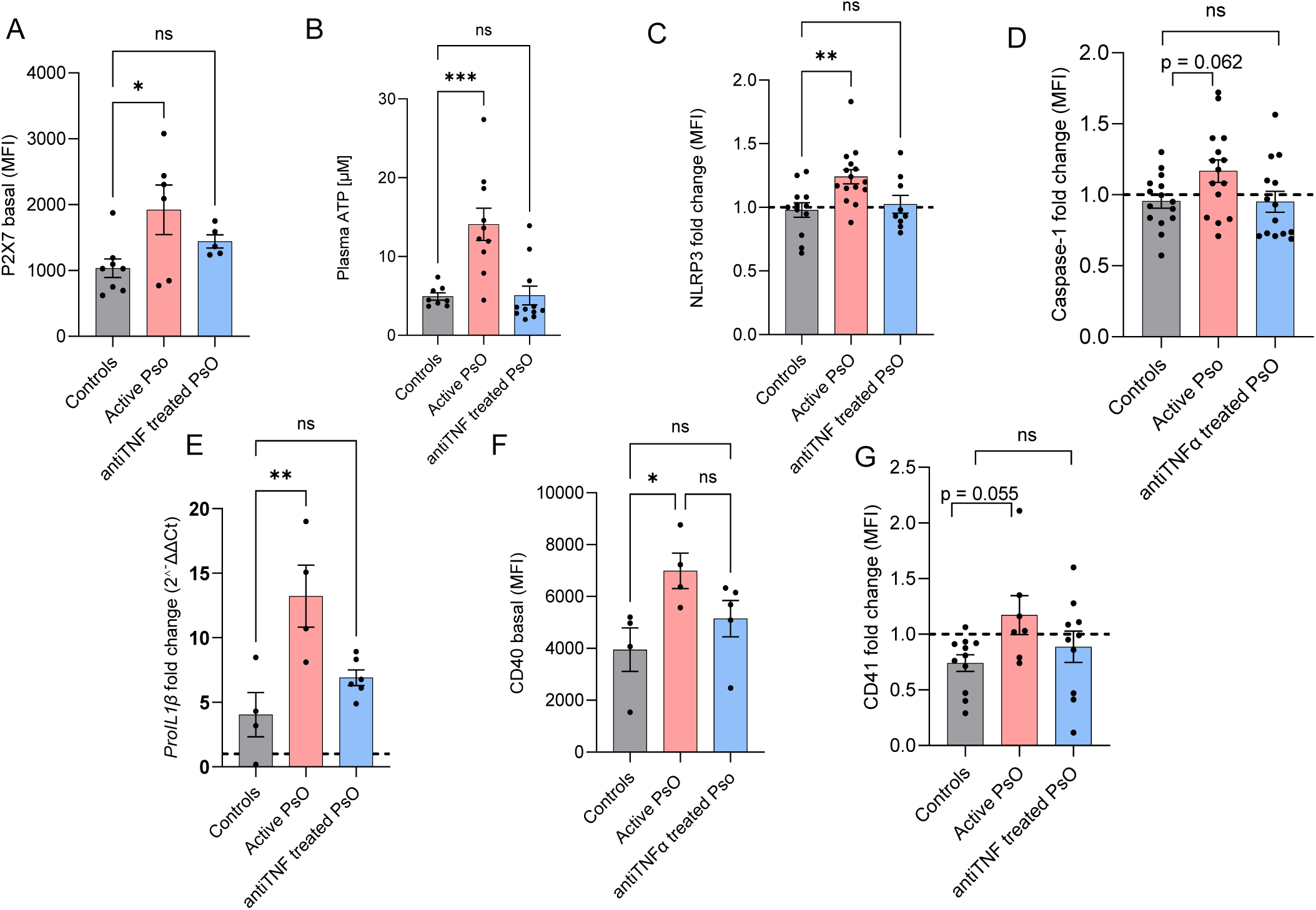
Anti-TNF- targeted treatment normalizes inflammasome signaling and platelet adhesion. (A) Ratio of NLRP3 (MFI) (B) caspase-1 (MFI) and (C) *pro-IL1B* (mRNA) expression at 37 °C versus room temperature for 10 minutes, as assessed by spectral flow cytometry and qPCR, respectively (n=4-17 per group). (D) Baseline P2X7 receptor surface expression (MFI) and (E) plasma ATP concentration (μM) measured by bioluminescence assay (n=5-11 per group). (F) Baseline CD40 expression (MFI) on classical monocytes, measured by spectral flow cytometry (n=4-5 per group). (G) Ratio of CD41 expression (MFI) on classical monocytes at 37 °C versus room temperature for 10 minutes, measured by spectral flow cytometry (n=8-10 per group). Data are presented as mean ±SEM. *p <0.05, ** p < 0.01, *** p<0.001. ns = non-significant. One-way ANOVA was used for comparisons involving more than two groups, and Šídák’s post hoc test was performed for multiple comparisons. Student’s t-test was used for comparisons between two groups.

In line with these results, patients receiving anti-TNF-α treatment, similar to healthy controls, did not exhibit increased NLRP3, caspase-1, or pro-*IL-1β* expression post-stimulation (Figure 6C-E). Our data suggest that anti-TNF-α therapy normalizes ATP signaling through P2X7R, thereby reducing inflammasome activation

### Anti-TNF treatment attenuates platelet aggregation to monocytes

Next, we investigated the effects of anti-TNF-α treatment on monocyte activation and platelet aggregation. At baseline, classical monocytes from anti-TNF-α-treated patients exhibited CD40 expression levels comparable to healthy controls (Figure 6F), indicating normalization of monocyte activation.

Upon stimulation, CD41 expression on monocytes in these patients was also similar to that of healthy individuals (Figure 6G), suggesting a restoration of normal platelet adhesion dynamics.

Collectively, these data highlight the immunomodulatory effects of anti-TNF therapy in psoriasis, demonstrating a restoration of immune cell function that likely contributes to reduced systemic inflammatory and thrombotic risk.

## DISCUSSION

Psoriasis has emerged as an independent risk factor for CVD ^4, 5^. The American College of Cardiology/American Heart Association currently includes psoriasis as a risk enhancer in the risk prediction score recommended to guide CVD prevention strategies ^29^. Although systemic inflammation in psoriasis is predicted to increase cardiovascular risk, the underlying cellular mechanisms have not yet been identified.

In this study, we demonstrate that in patients with active psoriasis, the NLRP3 inflammasome exhibits a lowered activation threshold compared to healthy controls, rendering immune cells hyperresponsive to minor perturbations in cellular homeostasis. This heightened sensitivity, or priming, was restricted to patients with active psoriasis and absent in those with inactive psoriasis. Thus, inactive psoriasis reflects a non-primed state consistent with clinical remission, aligning with evidence that cardiovascular disease risk correlates with the severity of the disease ^4, 30^.

We found a TNF-α-dependent increase in the expression of the cell surface receptor P2X7 on classical monocytes in psoriasis. The co-elevation of P2X7R and its ligand ATP in the plasma indicates a persistent systemic stress response, which likely contributes to the heightened systemic inflammation and increased cardiovascular risk in active psoriasis.

Recent studies demonstrated that P2X7 receptors play an important role in a variety of cardiovascular diseases, including atherosclerosis ^31, 32^. Interestingly, the P2X7 receptor was previously found to be increased in both lesional and non-lesional psoriatic skin compared to control ^33^, as well as on circulating monocytes in rheumatoid arthritis ^34^.

Extracellular ATP has previously been shown to be elevated in patients with atherosclerosis ^35^, hypertension ^36^, and SLE ^37^, underscoring its relevance in chronic inflammatory conditions. Since we found that normal PBMCs and those from active psoriasis express comparable levels of ATP-hydrolyzing ectonucleotidases CD39 and CD73, the increased extracellular ATP in active psoriasis likely arises from increased secretion, rather than impaired degradation.

In support of our observations of increased response to mild physiological stimulation, recent evidence reveals that NLRP3 activation is not only driven by canonical pathogen- or damage-associated patterns, but also by disruptions in cytoplasmic homeostasis ^38^. This paradigm shift helps explain persistent inflammation under sterile conditions and implicates altered cellular homeostasis as a predominant driver of NLRP3 activation. Notably, environmental stressors such as temperature fluctuations can disrupt cellular homeostasis and increase mitochondrial electron transport, resulting in a burst of ROS production ^39^. Damaged mitochondria serve as both sources of ROS and enhanced ATP synthesis, with mitochondrial ROS serving as a direct activator of NLRP3 ^40^.

Our proteomic profiling reveals baseline enrichment of endoplasmic reticulum and mitochondrial translation pathways in active psoriasis, even in the absence of external stimulation, indicating a constitutive mitochondrial stress state that likely predisposes immune cells to heightened oxidative responses upon activation. Consistent with this, we observed robust enrichment in mitochondrial ATP synthesis and ROS-generating pathways following stimulation. The most enriched pathway at baseline identified by our proteomic analysis was ‘enrichment of monocyte extravasation’, an unexpected finding given the relatively low percentage of monocytes (4–10%) in the whole blood samples. This robustly demonstrates a biologically significant upregulation of mechanisms that facilitate monocyte migration from the circulation into tissues, reflecting an activated immune state. Notably, monocyte recruitment into the vessel wall is a key early event in atherosclerosis ^41^. Supporting proteomic data, we found increased expression of CD40, a costimulatory molecule that amplifies inflammatory signaling and drives atherosclerotic plaque development ^42^, on circulating monocytes from patients with active psoriasis. CD40 stimulation induces production of inflammatory cytokines, chemokines, and matrix metalloproteinases by monocytes, contributing to plaque instability ^43^. Upon stimulation, we identified enriched pathways indicative of increased platelet activity and oxidative stress. Platelets are key players in inflammation and have been shown to enhance NLRP3 inflammasome activation in human monocytes by licensing *NLRP3* transcription and promoting optimal IL-1β release^44^. Biomarkers for platelet activation have previously been associated with psoriasis, such as platelet volume and soluble P-selectin ^45^.

The substantial enrichment of platelet adhesion pathways suggests a potential mechanism by which monocytes amplify the inflammatory cascade in psoriasis, further linking systemic inflammation to vascular dysfunction. The attenuation of CD41 expression on monocytes upon inhibition of mitochondrial ROS supports its role in platelet aggregation and strengthens the suggested link between immune and coagulation systems^46, 47^.

Among the key inflammatory mediators involved in psoriasis, TNF-α stands out for its role in innate immune activation ^48^. TNF-α not only promotes insulin resistance and endothelial dysfunction, both present in psoriasis and key contributors to CVD risk ^2, 49^, but also regulates inflammatory pathways, including the transcription of the NLRP3 inflammasome ^50^. In addition, TNF-α inhibitors have been linked to a reduction in CV events in patients with psoriasis ^51^.

The normalized NLRP3, caspase-1, and pro-IL-1β response in patients receiving TNF-α inhibitors provides a mechanistic explanation for the proposed cardiovascular risk-reducing effect. Since patients receiving TNF-α inhibitors demonstrated plasma ATP levels and P2X7 surface expression comparable to those of healthy individuals, it is plausible that treatment with TNF-α inhibitors, by restoring normal metabolic and immune function ^52^, indirectly influences ATP metabolism, thereby normalizing inflammasome activity.

Importantly, we found that anti-TNF-α therapy normalized both monocyte activation and formation of monocyte-platelet aggregates, underscoring the central role of TNF-α in restoring immune homeostasis and providing a mechanistic basis for reduced cardiovascular risk in anti-TNF-α-treated patients.

Patients with psoriasis included in this study had mild to moderate disease. Patients with severe disease were excluded due to their higher prevalence of comorbidities associated with increased cardiovascular risk. Moreover, our patient groups showed no significant differences in BMI compared to healthy controls. This approach allowed us to assess cardiovascular risk attributable primarily to psoriasis itself. Interestingly, our findings of increased systemic stress were not reflected in the conventional laboratory markers of systemic inflammation, such as hsCRP and WBC-derived markers, which are known to correlate with the occurrence of cardiovascular risk^53^. The cross-sectional nature of our study does not allow us to draw conclusions about whether inflammasome activation is causative of monocyte-platelet activation or merely correlated with it. However, the results from our hypothesis-generating findings are consistent with findings by Wu et al.^47^, who demonstrated that inflammasome activation can trigger blood clotting in sepsis.

In summary, our findings provide insights into how mitochondrial stress and disrupted cellular homeostasis may contribute to the primed, hyper-responsive state of innate immune cells in active psoriasis. The enrichment of platelet adhesion pathways suggests a possible mechanism through which monocytes amplify the inflammatory response, linking systemic inflammation to vascular dysfunction. Our findings support a role for TNF-α in mediating crosstalk between psoriatic inflammation and cardiovascular disease risk and show that TNF-α inhibition may help reduce systemic inflammasome activity and inflammatory burden.

## ACKNOWLEDGEMENTS

We would like to acknowledge the Core Facility, Faculty of Medicine and Health Sciences, Department of Biomedical and Clinical Sciences, Linköping University, for assistance with flow cytometry and mass spectrometry. We would also like to thank Ms. Eva-Lott Nord Carlsson för excellent technical assistance.

---

**Supplementary table S1.** List of antibodies.

| **Extracellular markers** | **Conjugate** | **Company** | **Panel** |
| --- | --- | --- | --- |
| CD3 | PerCP-Cy5.5 | BD Biosciences | 1 |
| CD4 | AH7 | BD Biosciences | 1 & 2 |
| CD8 | BV570 | Nordic Biosite | 1 & 2 |
| CD11b | BV650 | BD Biosciences | 1 |
| CD14 | APC Fire 810 | Nordic Biosite | 1 & 2 |
| CD14 | APC H7 | BD Biosciences | 1 |
| C16 | PE-Cy7 | BD Biosciences | 1 & 2 |
| CD33 | BV421 | BD Biosciences | 2 |
| CD56 | BV786 | Nordic Biosite | 1 & 2 |
| CD64 | PerCP-Cy5.5 | BD Biosciences | 2 |
| CD86 | APC | Nordic Biosite | 2 |
| CD192 | PE | BD Biosciences | 2 |
| HLA-DR | BV605 | BD Biosciences | 2 |
| P2X7 | AF405 | R&D Biosystems | 1 & 2 |
| **Intracellular markers** |  |  |  |
| NLRP3 | PE | Biotechne | 1 |
| Caspase-1 | FAM-YVAD-FMK | ImmunoChemistry | 2 |
| TOPRO3 |  | Thermo Fisher Scientific | 2 |

**Figure S1.**
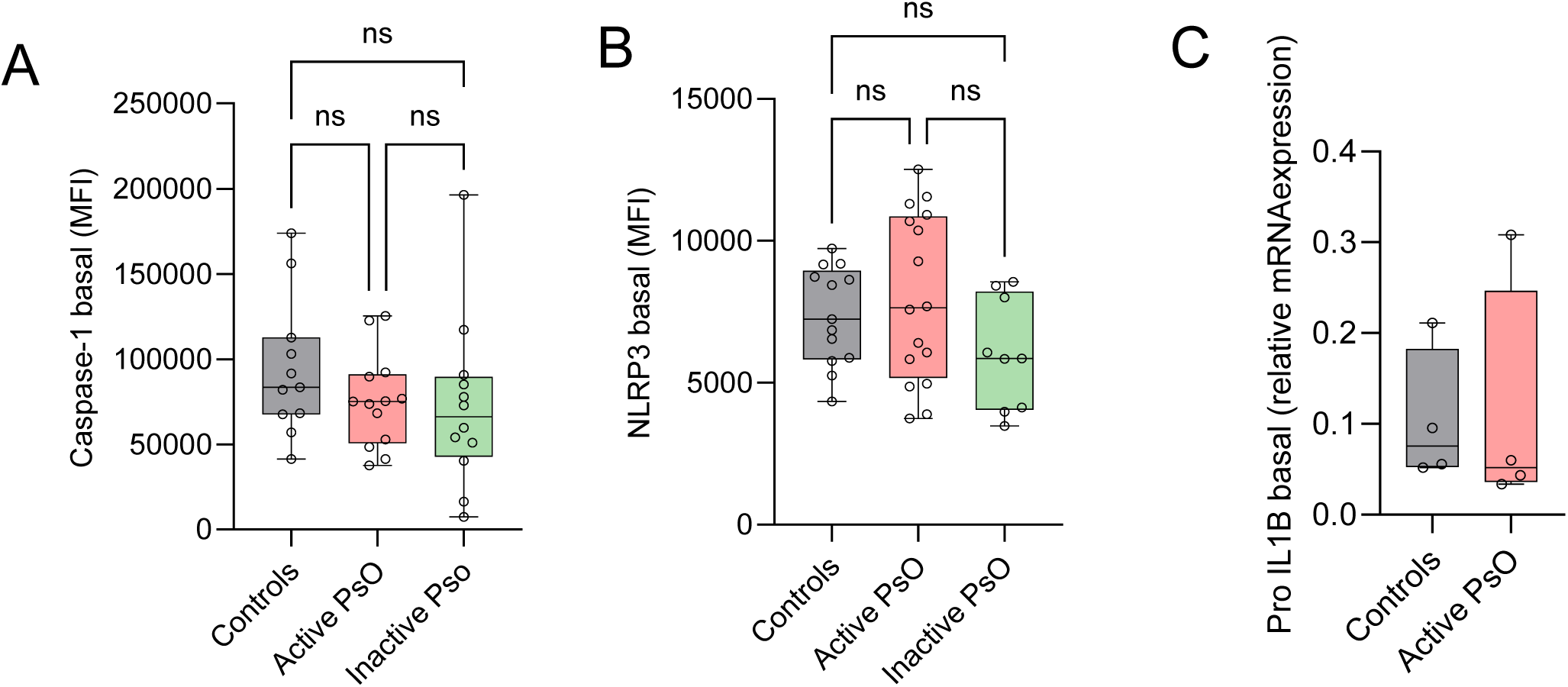
Basal expression of the inflammasome is similar between groups. Constitutive Caspase-1 (MFI), (B) NLRP3 (MFI) in controls, active and inactive psoriasis (PsO), and (C) *Pro-IL1β* mRNA in controls and active PsO. One-way ANOVA was used for comparisons involving more than two groups and Šídák’s post hoc test was performed for multiple comparisons, and Student’s t-test was used for comparisons between two groups. ns = non-significant

